# Evidence for a trade-off between glyphosate resistance and anti-grazer defence in green alga *Chlamydomonas reinhardtii*

**DOI:** 10.1101/2024.02.26.582141

**Authors:** Erika M Hansson, Dylan Z Childs, Andrew P Beckerman

## Abstract

1. The widespread and persistent use of herbicides has both selected for dramatically increased levels of herbicide resistance in the weed populations they were designed to control, and increased contamination of non-target ecosystems.
2. Experimental evolution using microbes offers the opportunity to explore basic evolutionary theory, including testing for the existence of intrinsic and extrinsic fitness trade-offs, connecting them to the patterns observed in both natural and agricultural populations.
3. We used green alga *Chlamydomonas reinhardtii* adapted to high and moderate levels of glyphosate to test for an extrinsic cost to glyphosate resistance in the form of a trade-off with clumping, the inducible anti-grazer defence deployed against gape-limited micrograzers. Through exposing the algae to freshwater rotifer *Brachionus calyciflorus* as well as their isolated info-chemicals, we test whether glyphosate resistance affects ability to deploy defences as well as ability to withstand grazing compared to control populations.
4. We find an increase in variation rather than uni-directional effect of glyphosate treatment, with treated populations exhibiting both lower and higher degree of anti-grazer defence compared to controls. Furthermore, our data suggests there are at least three different glyphosate resistant phenotypes present, with two conferring different extrinsic costs in the form of tradeoff with anti-grazer defence.
5. Our study indicates that extrinsic costs to glyphosate resistance may be context dependent in this system, possibly only being present at particular stages in the adaptation process or with particular resistance mechanisms.

## INTRODUCTION

Herbicides are used worldwide to protect crops by controlling weed populations. However, this widespread and persistent use also strongly selects for weeds to evolve resistance to herbicides, and herbicide resistant weeds is an increasingly costly problem, both economically and ecologically (Gaines *et al*., 2020; Powles & Yu, 2010). Furthermore, agricultural run-off means herbicides increasingly contaminate non-target ecosystems and assert a selective pressure for resistance on non-target species (Van Bruggen *et al*., 2018). At the heart of the problem for both agriculture and conservation is thus an evolutionary process, so characterising and managing the problem requires evolutionary thinking (Neve *et al*., 2009, 2014).

Classic evolutionary theory states that increased fitness in a new environment should trade off with fitness in the original environment, negative pleiotropic effects of the alleles involved cause a cost of adaptation (Purrington, 2000; Vila-Aiub *et al*., 2009b). Characterising potential fitness costs of herbicide resistance is important for understanding the resistance trait and its mechanism as well as how they constrain adaptation by preventing fixation of new alleles or maintaining polymorphisms (Futuyma & Moreno, 1988; Rainey *et al*., 2000). Fitness costs may be intrinsic or extrinsic, that is, either to do with processes within the organism, or arising primarily through the exposure to external stressors. Intrinsic mechanisms include diverted energy allocation from growth and reproduction to the new trait (Purrington, 2000; Vila-Aiub *et al*., 2009b), or involve disruption of normal cell function (Eschenburg *et al*., 2002; Fonseca *et al*., 2020; Funke *et al*., 2009; Healy-Fried *et al*., 2007). By contrast, extrinsic fitness costs only become apparent in certain environments, either due to interference with specific molecular processes or resource allocation costs affecting stress tolerance (Reznick *et al*., 2000; Strauss *et al*., 2002). This could reduce performance in extreme conditions (Ge *et al*., 2011; Vila-Aiub *et al*., 2013) or impact ecological interactions such as competition (Pedersen *et al*., 2007), disease response (Salzmann *et al*., 2008) or anti-herbivore defence (Gassmann & Futuyma, 2004; Gassmann, 2005).

Experimental evolution using microbes allows testing evolutionary theory in the lab to connect it to patterns observed in both natural and managed populations (Elena & Lenski, 2003). Green alga *Chlamydomonas reinhardtii* represents an important group of primary producers exposed to herbicide contamination (Van Bruggen *et al*., 2018) and has previously been used as a model system for experimental herbicide resistance evolution (Reboud *et al*., 2007; Reboud, 2002), including to characterise intrinsic costs of resistance (Hansson *et al*., 2024; Lagator *et al*., 2013a,b; Vogwill *et al*., 2012). Here we use *C. reinhardtii* populations adapted to lethal and sublethal concentrations of growth-inhibiting herbicide glyphosate to test for extrinsic fitness costs, specifically focusing on trade-offs with anti-grazing defence.

Many algae are grazed on by zooplankton that are limited by gape size, making inducible clump formation to increase apparent size a common, effective and possibly conserved defence trait (Bor-aas *et al*., 1998; de Carpentier *et al*., 2019; Fisher *et al*., 2016; Harris *et al*., 1989; Lürling & Beekman, 2006; Lürling & Van Donk, 1997; PanČiĆ & Kiørboe, 2018). *C. reinhardtii* is a useful model system for this as it clumps through two processes: palmelloid colony formation, where 4–16 non-motile cells share a cell wall (Harris *et al*., 1989; Lürling & Beekman, 2006; Nakamura *et al*., 1978), and flocculation, where a larger structure comprising up to thousands of cells is held together by an extra-cellular mucous matrix (de Carpentier *et al*., 2019; Fan *et al*., 2017; Harris *et al*., 1989; Lürling & Beekman, 2006; Sathe & Durand, 2016). Both appear to form through modified cell division (Harris *et al*., 1989; Lürling & Beekman, 2006; Nakamura *et al*., 1978), although flocculations may also aggregate from previously free-swimming algae in response to specific stressors (Sathe & Durand, 2016).

Short-term grazing and herbicide stress have been found to interact both synergistically and antagonistically in *C. reinhardtii* as well as be impacted by pre-exposure of either stressor (Fischer *et al*., 2012). However, it is necessary to consider the impact of adaptation to either stressor, not just acute stress (Baselga-Cervera *et al*., 2016). Without considering the potential trade-offs between the evolved traits, which may have knock-on effects on the whole food web, we cannot asses the long-term impact of herbicide contamination on aquatic ecosystems, nor can we make predications for how it may affect agriculture.

To test whether glyphosate resistance trades off with clumping in *C. reinhardtii*, we here compare the clumping response to freshwater rotifer *Brachionus calyciflorus* kairomones in glyphosate-adapted and susceptible strains of *C. reinhardtii*. Furthermore, we test whether the clearance rate of live rotifers depends on algal strain to evaluate whether any differences in clumping response translate to differences in effective defence.

## MATERIALS AND METHODS

### ALGAL STRAIN AND CULTURE CONDITIONS

16 *Chlamydomonas reinhardtii* strain Sager’s CC-1690 wild-type 21 populations were cultured in a continuous flow-through chemostats (following Hansson *et al*., 2022) in sterile Ebert algal medium (Ebert, 2013) at a dilution rate of 0.15/24h with a shared multichannel peristaltic pump (Watson-Marlow 205S/CA16). Prior to the experiment, the algae had been kept in static stock cultures for two years (Ebert algal medium, 25°C, 24h light), with batches being transferred to fresh medium fortnightly, then in continuous flow culture for two months under the experiment control conditions. Cultures were maintained at a volume of 380ml±5% while continuously mixed by bubbling. A light box provided white light from below with surrounding white light fluorescent bulbs giving a light level of 75 µmol m^-2^ s^-1^ 24h a day, at an internal temperature of 30°C.

### HERBICIDE TREATMENT AND EVOLUTION OF HERBICIDE RESISTANCE

The *C. reinhardtii* populations reached steady state (approximately 250 000 cells/ml) in control medium without glyphosate. From 14 days after inoculation, 5 chambers were exposed to 100 mg/L (lethal) and 5 were exposed to 50 mg/L glyphosate (sublethal) (PESTANAL®, analytical standard) Glyphosate inhibits growth by blocking the shikimate pathway (Steinrücken & Amrhein, 1980) and 100 mg/L completely inhibits population growth, whereas 50 mg/L results in a population growth reduction in batch culture (Hansson *et al*., 2024). 6 chambers continued to receive the control medium.

The full experimental design and effects glyphosate on population density and growth rate are presented in Hansson *et al*. (2024). Briefly, evolution of resistance to glyphosate was defined as a continuous decline in population density followed by a recovery to steady state. Population densities were monitored using flow cytometry (Beckman Coulter CytoFLEX). CytExpert (Beckman Coulter) was used to gate and count particles detected in the PerCP-A channel (Excitation: 488nm, Emission: 690/50 BP), a robust method for estimating algal density (Kadono *et al*., 2004) which was validated against manual haemocytometer counts. R (version 4.0.5, R Core Team, 2021) and package mgcv (Wood, 2011) were used to construct a hierarchical generalised additive model (hGAM) with the day-to-day growth rate for each population as the response and treatment and day as explanatory variables. While the pattern expected for resistance evolution was detected in response to the lethal dose, the model resulted in a flat fit for the sub-lethal populations comparable to the control populations. However, growth assays detected evolving resistance in the sublethal populations at a two week delay compared to the lethal populations.

The cultures were sampled for clumping and feeding assays 36 days after treatment start when all populations in both treatment groups were at steady state and were indicated to have evolved resistance, 17 days after the first evidence of glyphosate resistance in the lethal populations (Hansson *et al*., 2024).

### ROTIFER CULTURES AND ROTIFER WATER PREPARATION

Freshwater rotifer *Brachionus calyciflorus* resting eggs were obtained from Florida Aqua Farms, Inc. The rotifers were kept under standardised stock conditions in jars of 400 ml hard artificial pond water (“ASTM”, ASTM, 1989), with 24h 75 µmol m^-2^ s^-1^ warm, white light from above and below at 25°C. They were fed 50 µl of *Nannochloropsis* paste (Seahorsebreeder) daily.

Experimentally triggering a clumping response requires water that has had rotifers in it. This long established method works because metabolites from the rotifer grazers represent kairomone signals of grazer risk to the algae (Lürling & Van Donk, 1997). Rotifer water (“RW”) was prepared from the water of a 14 days old population where the population density averaged 100 rotifers/ml the 5 days before, which was filtered through 0.2 micron filters using a vacuum pump to prevent introduction of algae or bacteria from the rotifer culture (Lürling & Van Donk, 1997). The water was serially diluted with ASTM to the desired concentrations and stored frozen.

### DETERMINING THE EFFECT OF GLYPHOSATE RESISTANCE ON CLUMP FORMATION

To test the effect of glyphosate resistance on clumping response to *B. calyciflorus* kairomones in the absence of live rotifers, 4 technical replicates of 7500 algal cells from each *C. reinhardtii* population was washed with ASTM and placed in RW corresponding to a density of 100, 50 or 25 rotifers/ml or ASTM for controls. We measured clumping with a plate reader (Tecan Spark 10M Multimode Microplate Reader) at 96 hours after inoculation using confluence as a proxy. Confluence measures surface cover and gaps within and between cover in the well, with more cell aggregation as well as larger cells resulting in more coverage with fewer gaps and thus higher confluence. Pilot data using microscopy and flow cytometry indicate a strong positive relationship between confluence and clumping (ME Sorensen, pers. comm., April 2018). The populations were kept next to the chemostat populations to ensure as similar conditions as possible, in 30°C with 24h light provided by the light box and white light fluorescent bulbs at 75 µmol m^-2^ s^-1^.

### STATISTICAL ANALYSIS OF CLUMPING ASSAYS

All data were analysed using R (version 4.0.5, R Core Team, 2021). Linear mixed-effects models were fit with lme4 package (Bates *et al*., 2015) to test the effect of glyphosate and rotifer concentration as well as their interaction on confluence, with population chamber fit as a random effect with a varying intercept. The Anova() function from the car package (Fox & Weisberg, 2019) was used to test the significance of the fixed and random effects and confirmed through parametric bootstrapping using the pbkrtest package (Halekoh & Højsgaard, 2014). Effect sizes were estimated using the effectsize package (Ben-Shachar *et al*., 2020). The emmeans package (Lenth, 2022) was used for pairwise comparisons of the mixed-effects model estimated means. Specifically, a pairwise contrast between the confluence of controls at RW = 0 and RW = 100 established the baseline effect of RW. The average confluence for all glyphosate treated populations as RW = 0 and RW = 100 were contrasted to test for whether glyphosate effect varied by RW. Lastly, the confluence at RW = 100 was contrasted between controls and the lethal and sublethal dose populations separately to determine if either treatment reduced clumping.

### DETERMINING THE EFFECT OF GLYPHOSATE RESISTANCE ON CLEARANCE RATE

To the effectiveness of clumping as an anti-grazing defence and whether it was affected by glyphosate resistance, 6 technical replicates of 2 × 10^5^ cells from each *C. reinhardtii* population were washed with and suspended either in 5 ml of hard ASTM water with 40 adult pregnant *B. calyciflorus* females (n = 3) or without rotifers (controls, n = 3). The presence of live rotifers was confirmed and a sample of 400µl was removed at 0, 24, 48 and 72h to measure cell density using flow cytometry (Beckman Coulter CytoFLEX).

### STATISTICAL ANALYSIS OF FEEDING ASSAYS

R packages flowCore and ggcyto (Ellis *et al*., 2020) were used to determine population density by gating and count events detected in the PerCP-A channel (Excitation: 488nm, Emission: 690/50 BP) to determine population density. Rotifer clearance rate (CR) between the 0 and 72 hour timepoints was calculated as *CR* = *log*[(*Y* _t_ *− Y* _t-1_)*/*Δ*t*] *− log*[(*N* _t_ *− N* _t-1_)*/*Δ*t* where *N* is the population density in the absence of rotifers, *Y* is the population density in the presence of rotifers, and Δ*t* is the time elapsed between timepoints.

Linear mixed-effects models were fit with the lme4 package (Bates *et al*., 2015) with the absolute values of clearance rate as the response and glyphosate treatment line as a fixed effect, with population chamber fit as a random effect with a varying intercept. Bartlett’s test was used to test the effect of glyphosate treatment line on variation in the clearance rate, with clearance rates square root-transformed to meet the test’s normality assumption.

To confirm the expected negative relationship between confluence and clearance rate and identify outliers, linear regression was used to test the relationship between the mean confluence for each population at RW = 100 rotifers/ml and the clearance rate.

## RESULTS

### GLYPHOSATE TREATMENT EFFECT ON CLUMPING RESPONSE VARIES BY POPULATION

RW concentration had a strong effect on confluence (*χ*^2^ = 131.7, DF = 2, p < 0.001, *η*^2^ = 0.4), with considerably higher confluence at all rotifer concentrations compared to the controls for all *C. reinhardtii* populations but two (Figure 1, Table 1). Although the interaction between glyphosate treatment and RW concentration was not significant (interaction RW*Glyphosate *χ*^2^ = 9.9, DF = 6, p = 0.1, *η*^2^ = 0.06), the pairwise contrast estimated a smaller effect of RW in the glyphosate treated populations (Table 1), suggesting that the evolution of glyphosate resistance on average reduces clumping. However, no impact of glyphosate was found within RW treatment (*χ*^2^ = 1.1, DF = 2, p = 0.6, *η*^2^ = 0.08), with pairwise comparisons between controls and either herbicide treatment at the highest RW concentration showing no significant effect (Table 1).

**Table 1:** Selected pairwise contrasts of linear mixed-effects model fixed effect term means as estimated by the emmeans package for R. RW signifies RW concentration, H signifies glyphosate concentration.

| Contrast | estimate | SE | df | t-ratio | p-value |
| --- | --- | --- | --- | --- | --- |
| RW0H0–RW100H0 | -28.4 | 3.7 | 167.0 | -7.7 | < 0.001 |
| RW0meanH50H100–RW100meanH50H100 | -19.5 | 2.9 | 167.0 | -6.8 | < 0.001 |
| RW100H0–RW100H50 | 13.8 | 8.7 | 17.9 | 1.6 | 0.1 |
| RW100H0–RW100H100 | 3.2 | 8.7 | 17.9 | 0.4 | 0.7 |

**Figure 1:**
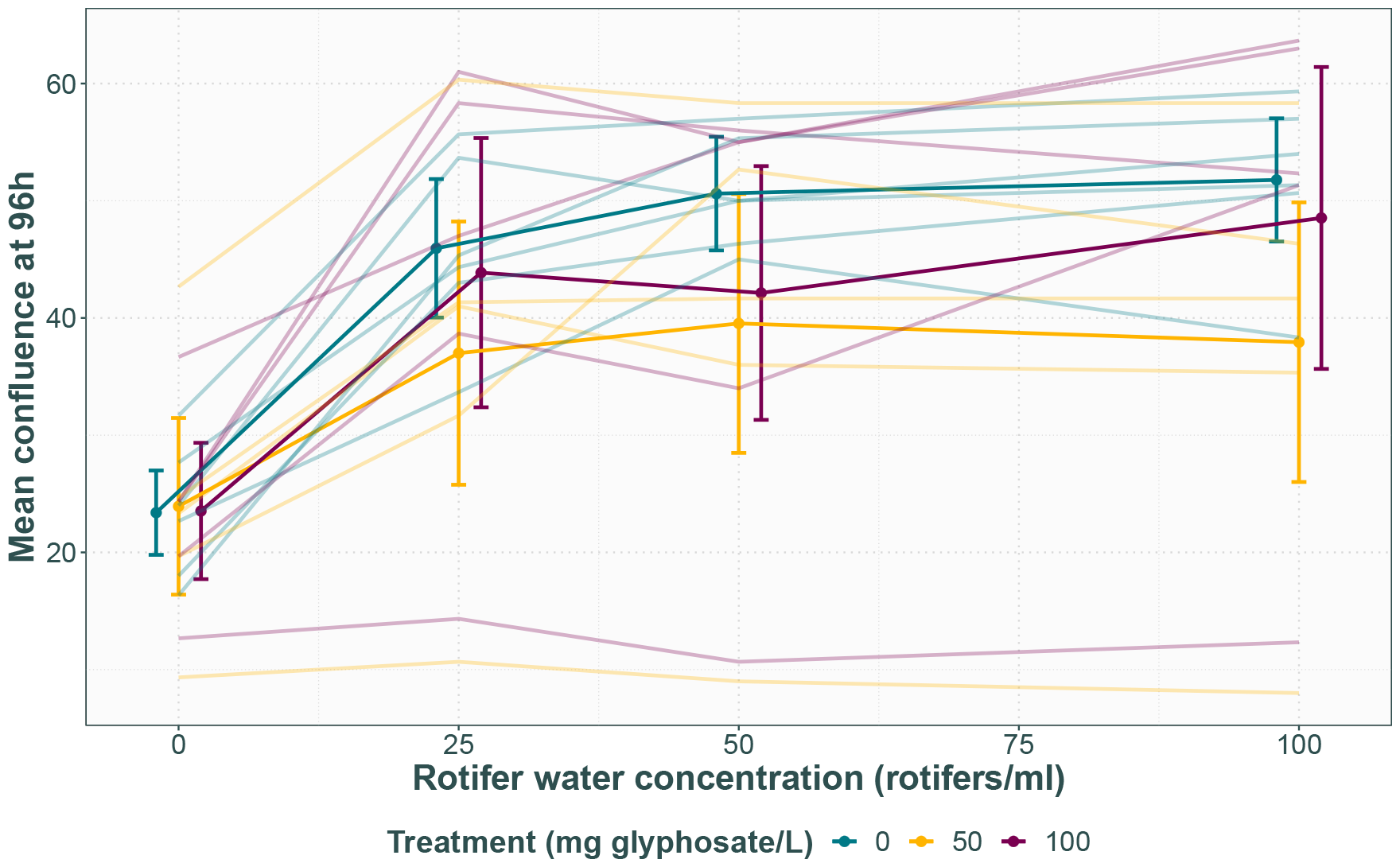
Mean confluence after 96 hours for each glyphosate treatment line in response to each rotifer water treatment. Opaque points with 95% CI represent the average for each glyphosate treatment, with transparent lines representing each replicate population.

There was also significant among population variation, with population replicate as a random effect having a highly significant effect (*χ*^2^ = 98.3, DF = 1, p < 0.001). Two populations, one from the lethal and one from the sublethal treatment, show consistently low confluence in the presence and absence of rotifers, suggesting they are not responding to the kairomones and possibly have a reduced growth rate.

### GLYPHOSATE TREATMENT EFFECT ON ROTIFER CLEARANCE RATE VARIES BY POPULATION

Glyphosate treatment did not have an effect on mean clearance rate (*χ*^2^ = 0.6, DF = 2, p = 0.7, *η*^2^ = 0.05), but it did have an effect on its variance (K^2^ = 18.1, DF = 2, p-value < 0.001), with both higher and lower clearance rates experienced by the glyphosate treated populations compared to the controls (Figure 2). Biological replicates from each population cluster together with similar clearance rate levels, and likelihood ratio testing of the random effect found a highly significant effect of population chamber (*χ*^2^ = 67.0, DF = 1, p < 0.001). This suggests that the increase invariance is due to each population representing an individual evolutionary trajectory and exhibiting different evolutionary strategies rather than an increase in noise.

**Figure 2:**
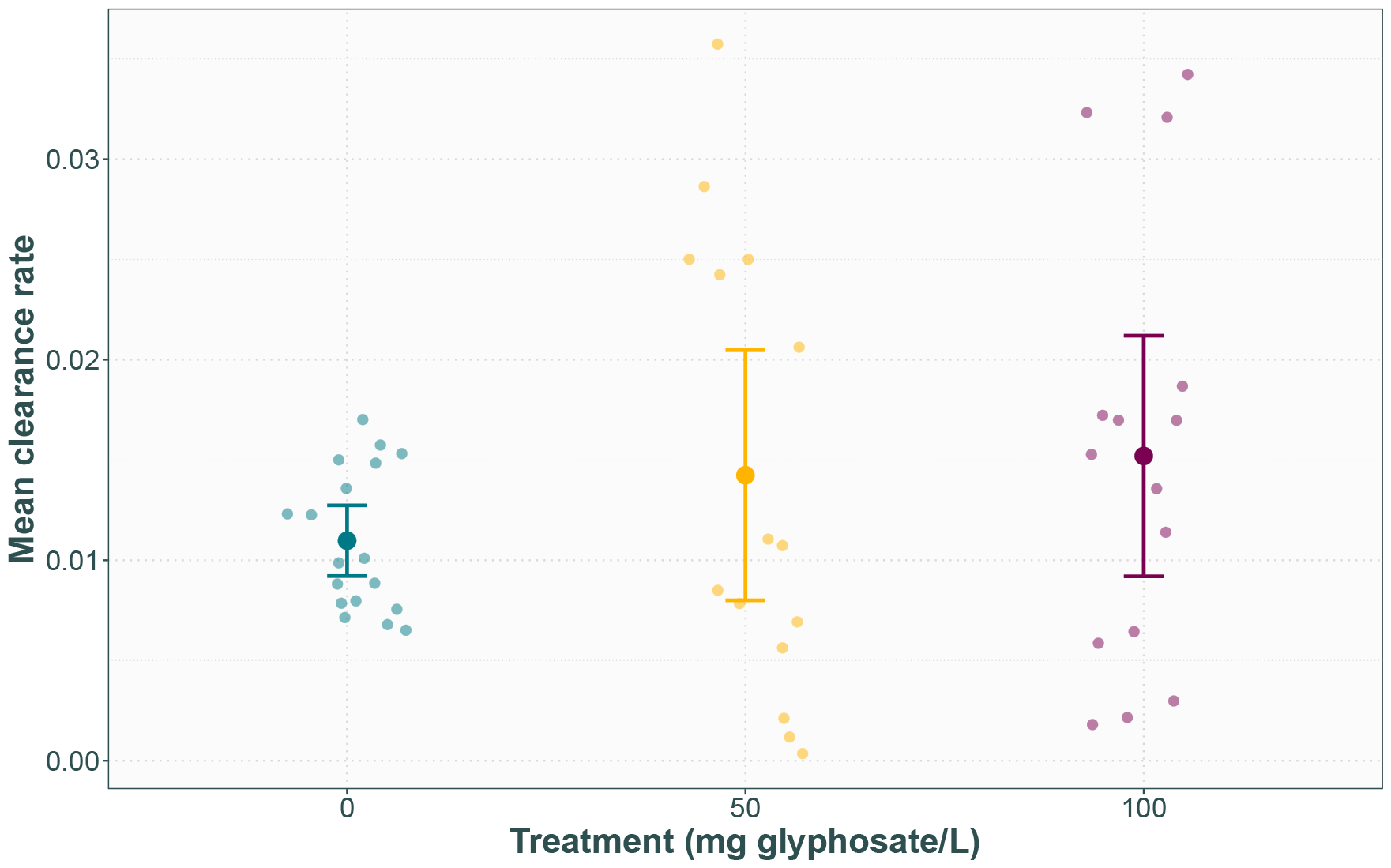
Mean clearance rate by *B. calyciflorus* feeding on algae from each glyphosate treatment line. Opaque points with 95% CI represent glyphosate treatment averages, and transparent points each assay replicate.

### HIGH CONFLUENCE CORRELATES WITH LOW CLEARANCE RATE IN MOST, BUT NOT ALL, POPULATIONS

The negative relationship between confluence and clearance rate (Figure 3) was found to be marginally significant, but with large effect size (F = 3.5, DF = 1,14, p = 0.08, *η*^2^ = 0.2), indicating that in most cases, confluence level indicates an adequate defence against grazing. The two outlier populations with lower confluence levels (A, B) do indeed experience higher clearance rates as expected. However, a third outlier population (C) from the sublethal treatment with high confluence similar to the controls also experienced a higher clearance rate, suggesting a separate mechanism is involved.

**Figure 3:**
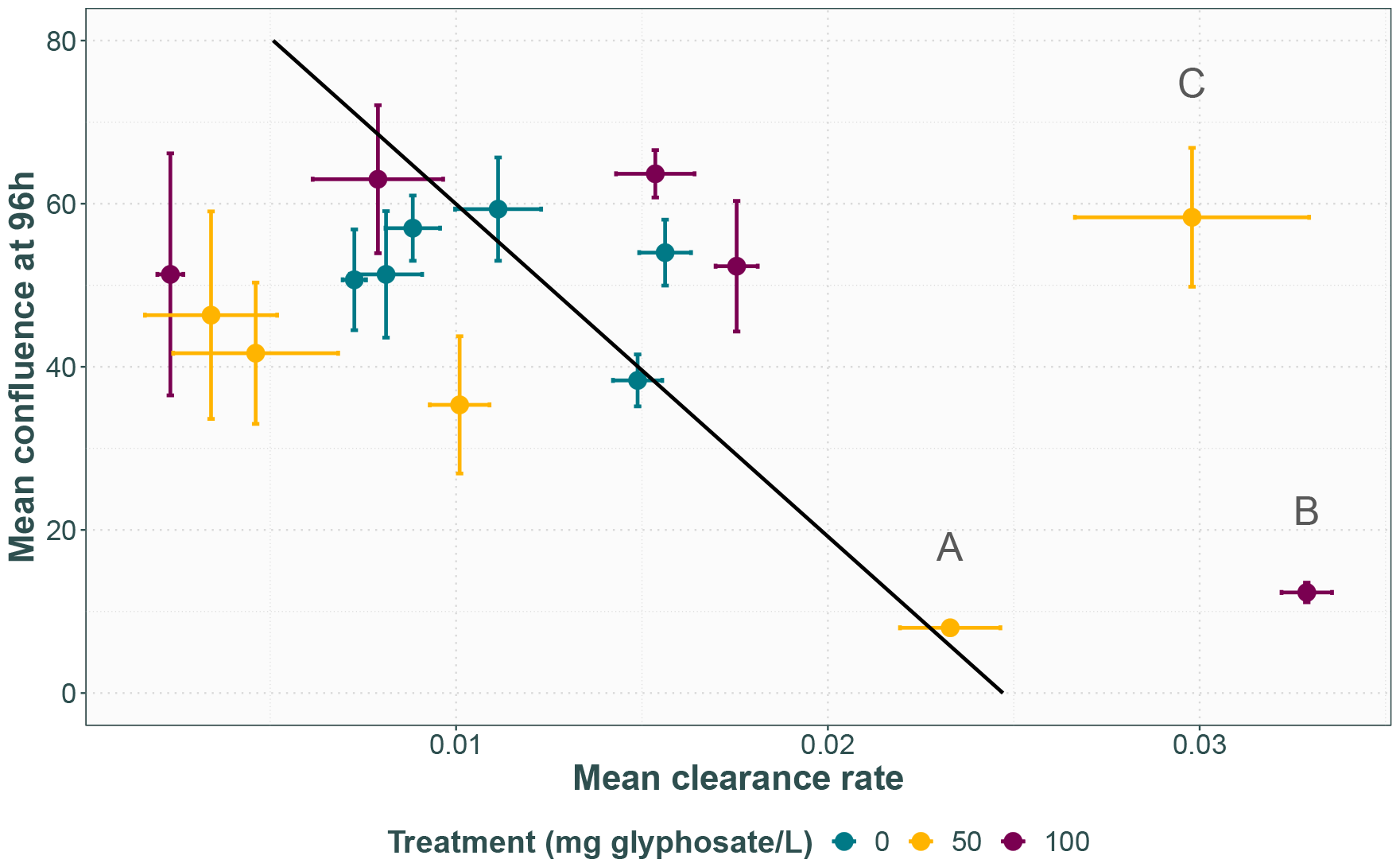
Mean confluence at 96h in the highest RW concentration plotted against mean clearance rate by *B. calyciflorus* for each given population with error bars for standard error. Regression line in black with equation *y* = 0.0247092 − 0.0002454*x*. Three outlier populations of interest have been labelled with letters.

## DISCUSSION

To predict the effects of herbicide resistance on agriculture and non-target ecosystems, we need to understand its evolutionary dynamics. This includes the fitness consequences of resistance in different ecological contexts as costs may be intrinsic or extrinsic. Model organisms such as *C. reinhardtii* provide the opportunity to test the evolutionary theory under controlled lab conditions with huge population sizes and fast generation times, giving insight both to weed science and evolutionary biology. We here focused on extrinsic costs by testing the ability of *C. reinhardtii* populations adapted to lethal and sublethal concentrations of glyphosate to deploy anti-grazer defences and withstand grazing by rotifer *B. calyciflorus*.

### ADAPTATION TO GLYPHOSATE INCREASES VARIATION IN ABILITY TO DEPLOY ANTI-GRAZER DEFENCE

The effect of glyphosate resistance on clumping and ability to withstand grazing by live rotifers is not consistent among populations, instead resulting in increased variation in defence deployment and efficacy compared to controls. Notably two populations, one lethal and one sublethal treatment, lost their ability to clump in response to rotifer kairomones. This was associated with an increased clearance rate by live rotifers. The other glyphosate treated populations clumped at the same level as the controls, but the clearance rates experienced were both lower and much higher.

This is consistent with each population representing a separate evolutionary trajectory, with different adaptive responses to the glyphosate treatment based on standing genetic variation and chance. Three distinct patterns emerge in performance by the glyphosate treated lines here: 1) performance same as controls, suggesting no trade-off between the glyphosate resistance trait and anti-grazer defence, 2) reduced ability to both clump and withstand grazing, indicating a trade-off with glyphosate resistance, and 3) ability to clump same as controls but reduced ability to withstand grazing, indicating a different trade-off with glyphosate resistance. This suggests there are at least three different dominant resistance mechanisms in the populations tested, two of which confer an extrinsic cost and at least one that does not.

### IMPLICATIONS FOR RESISTANCE AND TRADE-OFF MECHANISMS

The populations comprise several, coexisting lineages (Gresham & Hong, 2014; Maharjan *et al*., 2012) rather than isolated clones, meaning the assay results represent the averages of all phenotypes present rather than measuring any genotype exactly. Given the short time frame of adaptation, the underlying genotypes most likely 1) involve single substitutions to the target enzyme or up and down regulation of relevant metabolic pathways rather than resistance mechanisms dependent on sequential mutations (Gaines *et al*., 2020; Powles & Yu, 2010), 2) are still far away from the fitness optimum, but some may be further away than others (Gresham & Hong, 2014), and 3) owe their changes in performance to pleiotropy of the glyphosate resistance trait rather than neutral mutation accumulation.

A glyphosate resistant genotype resulting in both reduced clumping and defence could be based in limited resource availability (Purrington, 2000; Vila-Aiub *et al*., 2009a). Clumping may reduce nutrient uptake, both due to the extra-cellular mucous matrix hindering diffusion of vital nutrients (Becks *et al*., 2012) and due to larger cell sizes generally correlating with lowered metabolic activity (Marañón, 2015; Raven & Kubler, 2002), but this cost often only becomes apparent under nutrient limiting conditions (Becks *et al*., 2012; PanČiĆ & Kiørboe, 2018). Glyphosate blocks carbon metabolism, and single nucleotide substitutions conferring resistance by changing the target enzyme structure generally also reduce binding affinity for the intended substrates (Eschenburg *et al*., 2002; Fonseca *et al*., 2020; Funke *et al*., 2009; Healy-Fried *et al*., 2007), which leads to impaired cell functioning and affects resource availability. Furthermore, both flocculation and palmelloid formation in *C. reinhardtii* in response to *B. calyciflorus* is linked to mitosis (Harris *et al*., 1989; Lürling & Beekman, 2006), so any negatively pleiotropic effects on growth may also impair clumping ability, resulting in a negative feedback loop where lack of growth impairs clumping and the clumping that does occur further slows down growth due to reduced nutrient uptake.

By contrast, a genotype that still clumps in response to grazer kairomones, yet is less effective at withstanding the actual grazing, suggests that the clump formation is less robust. This could be due to lacking resources for production of the adhesive mucus, leading to increased disassociation when disturbed. Alternatively, part of the anti-grazer defence could be unrelated to colony formation and instead involve cell wall structure (Van Donk *et al*., 1997), cell hardness (DeMott, 1995) or the mucous envelope surrounding the cell (Porter, 1975) allowing some cells to pass through the rotifer gut without being digested. If instead this is trading off against the glyphosate resistance trait rather than colony formation, we would expect the pattern seen for the third outlier population (C). This trade-off could also be caused by limits on resource allocation, or perhaps changes to the cell wall or cell structure for a resistance mechanism like reduced absorption into the cell, although this is uncommon and tends to be accompanied by other mechanisms (Gaines *et al*., 2020; Nandula *et al*., 2013). Furthermore, lipid peroxidation of the cellular membranes from increased reactive oxygen species production is a known secondary glyphosate effect (Ahsan *et al*., 2008; Gomes & Juneau, 2016), evolved resistance against which could involve membrane composition.

An alternative explanation involving the secondary effects of glyphosate is that resistance mechanisms against the secondary effects of glyphosate are separate from the mechanisms lifting the primary shikimate pathway blockage, supported by glyphosate resistant weeds and genetically engineered resistant crops having been found to still be affected by the increased ROS production (Ahsan *et al*., 2008; Gomes & Juneau, 2016; Maroli *et al*., 2015, 2018). If so, the majority of the treated populations would in this scenario have evolved resistance mechanisms to withstand both, while the three outlier populations may have evolved resistance against the primary effects, but not the secondary. They would then still be suffering the effects of lipid peroxidation from chronic glyphosate exposure which could affect both growth and cell structure and thus potentially both clump formation and digestibility. This suggests that the stage of resistance evolution may influence the ability to respond adequately to additional stressors.

### BROADER IMPLICATIONS AND FUTURE RESEARCH

Our results suggest extrinsic fitness costs to herbicide resistance are not universal in this system, and may be dependent on timing, standing genetic variation and chance. Previous studies on the trade-offs between herbicide resistance and anti-grazer defence specifically concern a single plant– herbivore–herbicide interaction (*Amaranthus hybridus*–*Disonycha glabrata*–triazine, Gassmann & Futuyma, 2004; Gassmann, 2005). Furthermore, the current understanding of the general effects of herbicides and herbicide resistance on consumer-resource interactions is overall limited, despite the potential of herbicides for both strengthening and disrupting ecological interactions with dramatic consequences for whole communities and ecosystems (Iriart *et al*., 2021). This knowledge gap is particularly pertinent for algae as globally vital primary producers with the potential for rapid adaptation (Baselga-Cervera *et al*., 2016).

Ultimately, it is necessary to consider adaptation to stressors as a process as well as maintain a whole ecosystem perspective. This study contributes one piece of the puzzle for how anti-grazer clumping and glyphosate resistance in *C. reinhardtii* may shape evolution together at an early stage in adaptation. To determine the mechanism for the trade-off, microscopy analysis could distinguish whether flocculation or palmelloid formation is primarily involved, and analysis of the extra-cellular mucus could determine whether its composition is affected (Roccuzzo *et al*., 2020). Future studies should investigate the outcome of simultaneous glyphosate and rotifer stress at different stages of the adaptation process to determine its effect on the evolutionary trajectory. Furthermore, glyphosate may have direct effects on *B. calyciflorus*, such as stimulating sexual reproduction (Xi & Feng, 2004), which could further change the interaction, if they were exposed concurrently.

## ACKNOWLEDGEMENTS

Thanks to Megan Sorensen and Holly Banks for testing out the clumping assay method. Thanks to Allison Blake and Lynsey Gregory for technical support and maintenance of algal stocks. Thanks to Aga Urbanek for rotifer maintenance advice.

## AUTHOR CONTRIBUTIONS

EMH, DZC and APB conceived the ideas and designed methodology; EMH collected and analysed the data as well as led the writing of the manuscript. All authors contributed critically to the drafts and gave final approval for publication.

## DATA AVAILABILITY STATEMENT

For the purpose of open access, the author has applied a Creative Commons Attribution (CC BY) licence to any Author Accepted Manuscript version arising. The data presented in this manuscript, along with relevant code, is available at DOI: ZENODO

