## Supplementary materials 1 for "Evidence for a trade-off between glyphosate resistance and anti-grazer defence in green alga *Chlamydomonas reinhardtii*"

Erika M Hansson[1*], Dylan Z Childs[1], Andrew P Beckerman[1]
**1** School of Biosciences, The University of Sheffield, Sheffield, South Yorkshire, United Kingdom

*

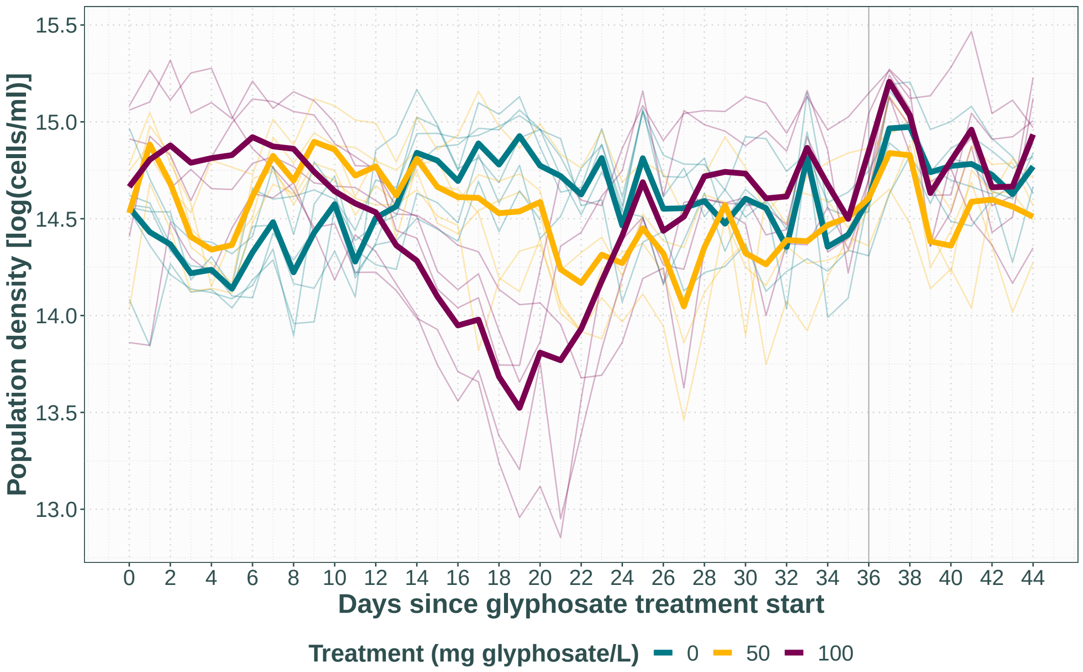


**Figure 1:** Data from (**Hansson2022intcost**). Average population density for each replicate population is shown in transparent thin lines, averages for each treatment are shown in thick opaque lines. The grey line at day 36 represents when samples for clumping and feeding assays were extracted.
